# Fight, Retreat, Repeat: Field observation of male-male agonistic behavior in the wood-feeding cockroach, *Panesthia angustipennis spadica* (Dictyoptera: Blattodea: Blaberidae)

**DOI:** 10.1101/2024.05.09.593021

**Authors:** Haruka Osaki, Tomohiro Nakazono, Kiyotaka Yabe, Mamoru Takata, Aram Mikaelyan

**Affiliations:** Department of Entomology and Plant Pathology, North Carolina State University, 100 Derieux Place, Raleigh, NC 27695, USA; Laboratory of Insect Ecology, Graduate School of Agriculture, Kyoto University, Kitashirakawa-Oiwake-cho, Sakyo-ku, Kyoto, 606-8502, Japan

**Keywords:** field observation, male-male agonistic behavior, *Panesthia*, *Salganea*, subsocial, wood-feeding cockroach

## Abstract

Conflict is one of the most critical factors affecting the behavior of animals related to their reproduction and survival, with aggressive interactions being central to acquiring resources or mating partners. This phenomenon is more common among males than females, impacting reproduction strategy and beginning of biparental care. Investigating such interactions in species closely related to social species can be illuminating, offering valuable insights into the factors that influence the emergence and maintenance of more complex social behaviors. In this context, we present a field study of male-male agonistic behavior in the wood-feeding cockroach, *Panesthia angustipennis. Panesthia* is the closest genus to the subsocial genus *Salganea*, which is known for its biparental care. Our field observations reveal a characteristic behavior where one male pushes a rival away from a female. The victorious male repeatedly returns to a specific site near the female, suggesting a strategy to minimize unnecessary conflict or protect the female. This behavior provides insights into the potential evolutionary strategies that may have evolved in the common ancestor shared by *Salganea* and *Panesthia*. Notably, the displaced males persistently reengage, highlighting the high resource value attributed to females and the consequential intensity of male competition. This study not only sheds light on the aggressive and pacifist tendencies in *P. angustipennis* but also contributes to understanding the evolutionary development of social structures in *Salganea*. Further experimental investigations into the aggressive behaviors of *P. angustipennis* will enhance our comprehension of the factors shaping the evolution of sociality in these species.

## Introduction

Male-male agonistic behavior is a universal phenomenon reported across diverse organisms (Pandolfi et al. 2021). After this aggressive behavior, the winner usually obtains the chance to courtship or is chosen by females. It affects mating decisions, impacting individual fitness and overall population dynamics (Andersson and Iwasa 1996). Such agonistic behaviors over mates shape the evolutionary course of species and their strategies, including sociality. Even after mating, it is well known that male-male agonistic behavior occurs to defend the females from other males (Suzuki 1999; Kvarnemo 2005). Protecting females and offspring by males is one of the proximate factors in the evolution of biparental care (Royle et al. 2016). Therefore, male-male agonistic behavior in closely related non-social species can offer valuable insights into the evolutionary drivers in social insects.

In this regard, *Panesthia angustipennis*, a wood-feeding cockroach, presents a unique case study. *Panesthia* is one of the closest genera to the subsocial wood-feeding cockroach, genus *Salganea* (Djernæs and Varadinova 2020), which is known for its biparental family structures (Maekawa et al. 2008) and shares many habits with *Salganea*. Despite the lack of documented male-male agonistic behavior in *Salganea*, the study of *P. angustipennis* offers remarkable insights. This species, active outside the log during the breeding season, tunnels into rotten logs (Asahina 1991) and exhibits ovoviviparity, with significant gestation periods. Given the skewed field sex ratio indicating shorter adult longevity in males, male-male agonistic behavior in *P. angustipennis* has been anticipated (Ito and Osawa 2019).

In this paper, we describe the male-male agonistic behavior of *P. angustipennis* as a species sharing the common ancestor with *Salganea*, presenting field observations that illuminate not only the behavioral dynamics but also its environmental context. By focusing on a species closely related to biparental cockroaches, our study offers fresh insights into the evolutionary underpinnings of aggressive behavior and its implications for obtaining sociality.

## Materials and Methods

### Field observation and collection

On August 26, 2022, we eventually observed two males engaged in a struggle on two logs in Kyoto, Japan (34°52’36.9”N, 135°51’59.3”E) at 7:21 p.m. The event was documented on video using a smartphone (iPhone SE; Apple, Cupertino, US) with headlights (GH-100RG; Gentos, Tokyo). The recording was terminated after 1:33 minutes into the struggle due to limitations imposed by the equipment. The two males, along with a female, were brought back to the laboratory, and the pronotum widths were measured. All three individuals were adults and had wings. The males were distinguished by their wings: Male X had torn wings, and Male Y had intact wings.

### Analysis

All behaviors were analyzed using the software Boris (version 7.9.7; (Friard and Gamba 2016)). We compared the observed behaviors with the following behaviors, which were the ones of another species in the same genus, *Panesthia cribrata*, reported in laboratory observation by O’Neill et al. (1987): (i) *Push*: lowering their pronota by pulling their heads down towards their legs. (ii) *Pulse*: extending and contracting the abdomen. (iii) *Block*: turning away from its opponent and lowering the side of the body or tip of the abdomen facing its opponent. (iv) *Submission*: standing still, abdomen downturned at the tip, and antennae motionless. If the behaviors not reported were observed, we record them as new behaviors. Statistical analysis was conducted with R version 4.3.2 (R Core Team 2023). We conducted the Friedman test to compare the behavioral counts demonstrated by each male across treatments. We were also interested in the difference between the two males in the number of counts of the behaviors when data was summed across treatments. Fisher’s exact test was used to examine the independence of those behaviors.

## Results

### Pronotum widths

Size measurements revealed that the pronotum widths of the female, Male X and Male Y, measured at 11.5 mm, 13.5 mm, and 13.1 mm, respectively.

### Field observation

The whole video observation is available in Online Resource 1. Among the four behaviors that had been reported, we observed two: *push* (Fig. 1a) and *block*. The newly observed behaviors are as follows:

**Fig. 1.**
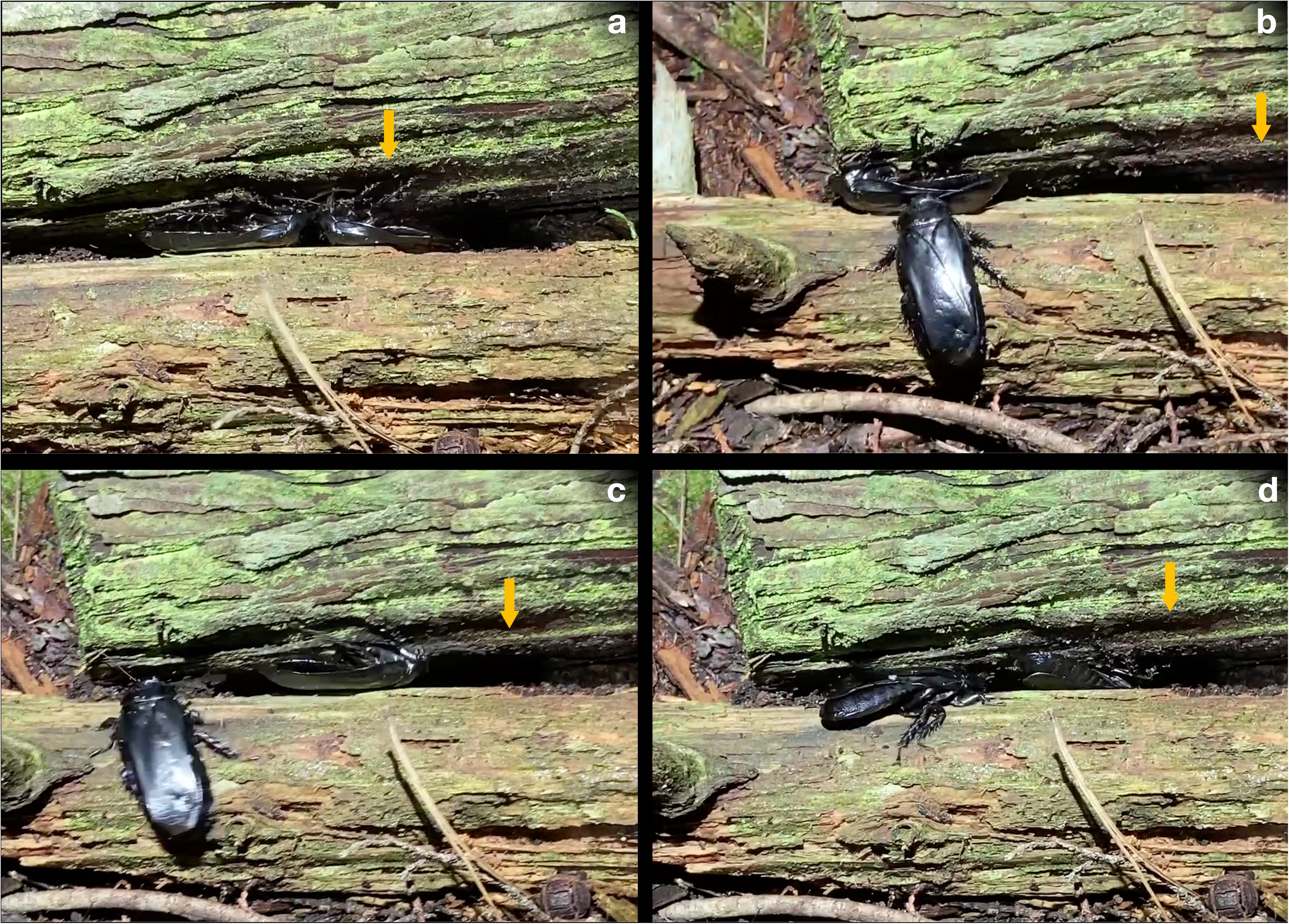
The behaviors observed during the male-male agonistic behavior in the field. The arrow indicates the location of the home position. The arrow seems to move because of the camera work. However, these four arrows indicate the same place. (a) The two males are *pushing* each other. (b) The lower male is *retreating*. (c) The right male is *returning*. (d) The left male is *pursuing* the opponent.

#### Retreat

A reaction in one male to being *pushed* or approached by another male, in which the male backs away without pivoting, often by evading any bout with the opponent (Fig. 1b).

#### Return

The male pivots and withdraws from the other male after a bout, consistently to the same location in the arena defined as “home position”, which is situated directly above the female’s location (Fig. 1c).

#### Pursue

One male pursues the other male who is *returning* (Fig. 1d).

#### Stop

A male ceases movement altogether.

The individual interactions observed as part of this agonistic behavior are as follows:

1. Initiation of the agonism with one male *pushing* the other.
2. A reciprocal *pushing* interaction, where both males engage in *pushing*.
3. Male X disengages and *returns* to the home position location within the arena.
4. Male Y *pursues* Male X again.
5. The sequence of interactions repeats through steps 1–4.

As shown in Figure 2, Male X demonstrated both *block* and *return* behaviors, which were not exhibited by Male Y. Specifically, Male X displayed a high frequency of *push* and a moderate occurrence of *return*, while Male Y primarily showed *retreat*.

**Fig. 2.**
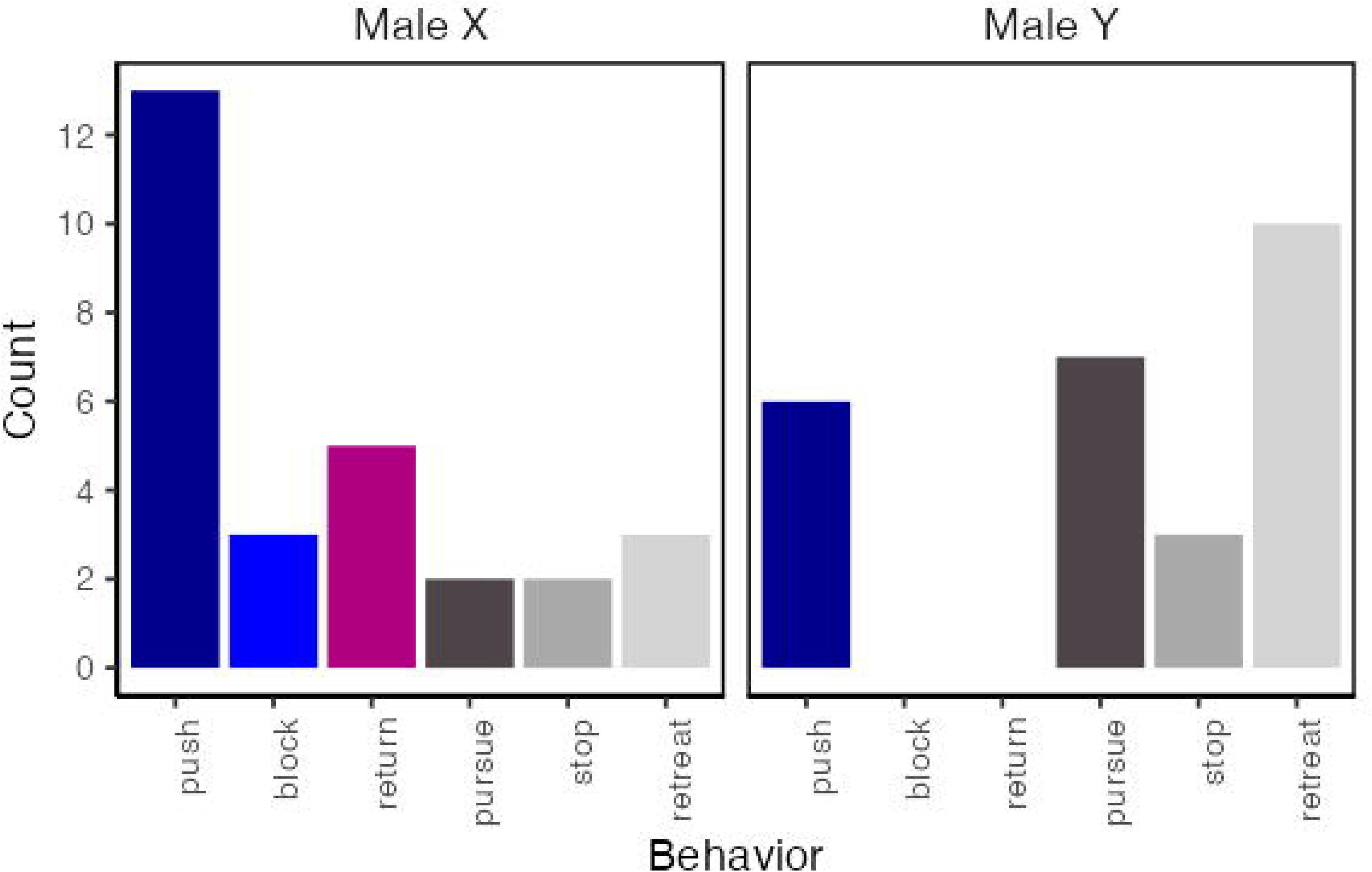
The behavior count recorded for Male X and Y in field observation.

This pattern of bout was observed to repeat 16 times, predominantly initiated by Male X (13 times), and less frequently, Male Y (3 times). Specific to Male X were the *return* and *block* behaviors; the *return* was documented five times, which included *return* once each at 3, 7, and 15 cm from home position, and twice at 10 cm from home position. The male *returned* at each place and went to home position. *Block* occurred three times, with two instances at 1 cm and one at 2 cm from home position. The video did not capture her because the female remained stationary under the gap between the two logs.

None of the males’ bouts involved the female. After observation, we found her under the log. The female remained absent from the fighting.

## Discussion

Our observations of male-male agonistic behavior in *Panesthia angustipennis* have unveiled a combination of aggressive and avoidance strategies that likely play a significant role in the evolutionary trajectory of social behavior in blaberid cockroaches. The repeated *push* and *return* behavior, particularly the *return* of the male to a home position near the female, suggests a delicate balance between aggression and avoidance. This balance may reflect an evolutionary strategy aimed at maximizing reproductive success while minimizing the costs associated with prolonged conflict, such as overexertion or injury.

Importantly, the male *returned* without completely subduing the invader. The range defended by the male is approximately twice the length of its body (around 10 cm), indicating that males preferentially defend a localized area around the female. This behavior highlights that the guarding male not only has a predisposition to minimizing conflict, but prioritizes proximity to females over the complete removal of its rival. Given that *Panesthia* is the closest genus to *Salganea*, which has subsociality including biparental care throughout its life, *returning* to the female is consistent with the hypothesis that the protection by males contributed to the evolution of biparental care (Royle et al. 2016). In addition, it is reasonable that the common ancestor has the nature to avoid unnecessary fighting because it indicates avoiding the wound and can contribute to the male’s longevity.

A male who was *pushed* out of the home position would continue to fight. This male showed his persistence by repeating *pursue*. The repeated attempts to initiate struggles, even after expulsion, were another characteristic of the agonistic behavior in *P. angustipennis* other than *return*. Generally, avoiding more fighting than necessary can reduce the cost of injury and maintain higher expectations of reproductive success (Smith and Price 1973; Arnott and Elwood 2008). The aggressive behavior is demonstrated to correlate to resource value (Enquist and Leimar 1987; Liu and Hao 2019). One of the possible reasons is the relatively short lifespan of *P. angustipennis* males, which are considered to have higher mortality than females (Ito and Osawa 2019). Because of shorter longevity, males might have limited opportunities to find females in their reproductive period. Another possible reason for repetition is their body sizes. The persistent struggle between the males could be attributed to the similarity in their body sizes, which likely complicates the establishment of dominance. This phenomenon has been noted in honeyeaters and spiders, wherein small differences in body size can intensify or prolong the struggles (Austad 1983; Kojima and Lin 2017).

The *block*, observed only in Male X as well as *push* and particularly within close proximity to the home position, highlights its tactic using two different types of defense. The block has been described as an effective defense only in a small gallery by laboratory observation (O’Neill et al. 1987). However, we observed *blocks* outside galleries despite less effectiveness. The *block* requires no changing the position differently from the *push*, which requires turning the head to the other male. The male was considered to choose *block* as an instantaneous option in case the other male got too close toward the home position to *push* it.

They *pushed* each other using the pronotum, which displays sexual dimorphism with more pronounced horns in males. From our observation, they were thought to serve as a weapon in *P. angustipennis*. This physical characteristic is also shared by pronotums in other blaberid cockroaches, such as hissing cockroaches (Durrant et al. 2016), and further underscores their importance in sexual selection and conflict resolution. Notably, however, this sexual dimorphism is not common in the subsocial *Salganea* (Asahina 1988), which might reflect a divergence in behavior in *Salganea* to exhibit less aggressive traits than *Panesthia*.

There is a strong need for more controlled laboratory experiments to explore further the triggers and consequences of the behaviors we observed in the field. Although we tried to observe those behaviors using the same individuals also in the laboratory, we were unable to record the *return* that we observed in the field. Furthermore, Male Y did not engage in fighting and consistently escaped from Male X over three days (Online Resource 2). While the results provided interesting insights, there is room to reconsider the design of the laboratory observation, perhaps with more individuals.

Our study suggests that antagonistic interactions in the males of *P. angustipennis* are a calibrated balance of two key strategies: attack and avoidance. We hypothesize that this strategy of “minimum fighting” may have contributed to the evolution of subsociality in the closely related species *Salganea*. This observation enhances our understanding of the evolutionary pathways leading to sociality by highlighting the complex behaviors involved in mate competition and conflict resolution. By documenting these behaviors in a species closely related to subsocial cockroaches, our study provides valuable insights into the ancestral traits that may have facilitated the evolution of social behavior in insects.

## Supporting information

Online Resource 1 & 2

## Acknowledgment

We thank Dr. Eiiti Kasuya for the valuable discussion on this study, and Dr. Kenji Matsuura for providing a well-equipped environment that facilitated our laboratory experiments. This study was supported by the Japan Society for the Promotion of Science Grant-in-Aid for JSPS Fellows Grant Numbers JP22KJ1779 (HO) and KAKENHI Grant Numbers JP21K14863 (MT). The work by AM was supported by the National Institute of Food and Agriculture Hatch project accession number 1019324.

## Data availability statement

The data including digital video images and data for all analyses can be found in Supplementary information.

