## Supplementary material for "Fight, Retreat, Repeat: Field observation of male-male agonistic behavior in the wood-feeding cockroach, *Panesthia angustipennis spadica* (Dictyoptera: Blattodea: Blaberidae)": Online Resource 1 & 2

The video is available here: https://youtu.be/Y4UVyQPZ4gY

**Online Resource 2**

**Laboratory observations:**

Method

On the day following the collection, we conducted a series of experiments with the two males and the female. Our observations were recorded using a video camera (HC-VX992MS; Panasonic, Tokyo, Japan) in controlled arenas (W200 × D150 × H100 mm). The experimental procedures were as follows: (i) Introduction of Male Y into the arena with Male X and the female. (ii) Simultaneous introduction of both males without the female. (iii) Introduction of Male X into the arena with Male Y and the female. In treatments involving the female, (i and iii), a Petri dish (φ90 × 150 mm) with an entrance was used to ensure proximity between the males and the female at the beginning of the introduction. The female and male were entered the petri dish 10 minutes before introducing the other male. Although, in treatment (iii), the pair exited from the petri dish during the 10 minutes, the introduction was conducted because the female and male were close to each other. All video recordings were terminated one minute after the males had ceased their physical encounter. Following each test, the individuals were isolated for 24 hours in the rearing system for wood-feeding cockroaches (Osaki 2022).

Results

Recording times varied between observations: 27 minutes for treatment (i), 6 minutes for treatment (ii), and 59 minutes for treatment (iii). Notably, in all treatments, Male Y ran away from Male X, which involved the male running in the opposite direction of their fighting partners. This behavior was named *escape* and observed in Male Y in all treatments (Fig. S1).

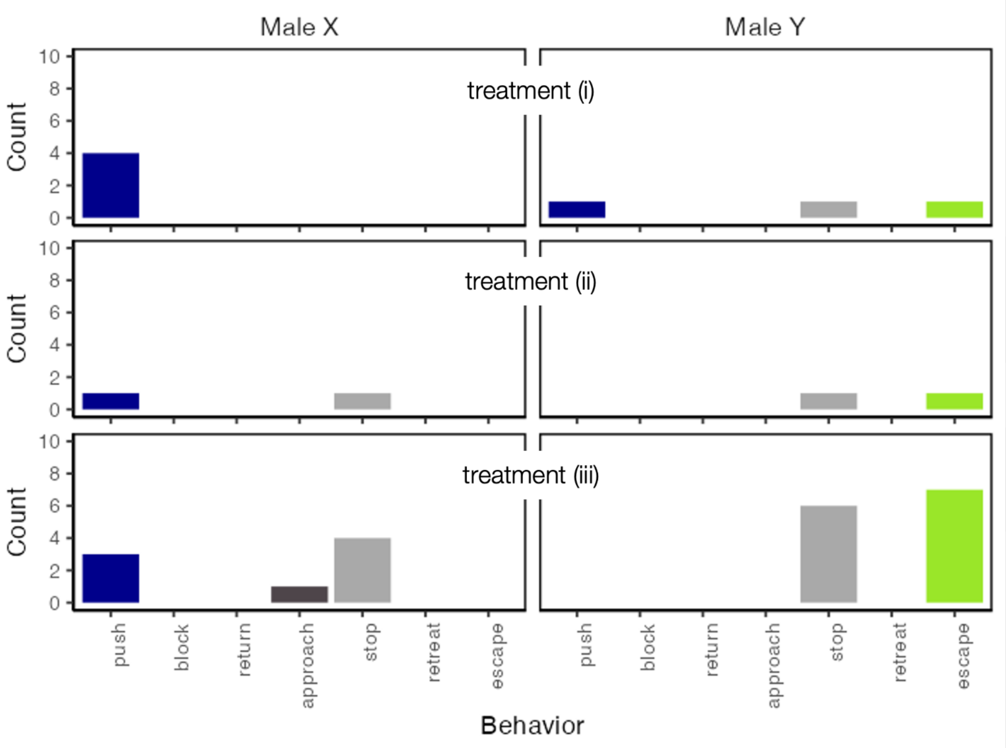


S1. Behavior counts in response to different treatments for Male X and Y in laboratory observation.

There were no significant differences observed in the number of times of behaviors demonstrated by each male across treatments (Friedman test, Friedman $\chi^2 = 4.3$, df = 2, p = 0.12). However, a significant difference was observed between the two males in the number of the *push* and *escape* when data was summed across treatments. Male X engaged more frequently in *pushes* while Male Y engaged more frequently in *escapes* (Fisher's exact test, $p < 0.05$).

The observed sequence of the agonistic behavior remained consistent, regardless of the female's presence.

1. Initiation to antennate by one male.
2. The initiating male *pushes* the other male.
3. Male Y executes an *escape*.
4. Male X, often moving with heightened activity, would antennate with Male Y.
5. The sequence of interactions repeats through steps 1–4.

Agonistic behavior was initiated upon antennating, leading to one male pushing the other. Regardless of which of the males initiated the interaction, Male Y consistently *escaped*. Even upon *escape*, when the struggle between the males was staged again, Male Y's response to Male X's *push* was always to *escape* rather than to *push*.

The female did not join the fighting. Throughout the observation of these struggles, the female exhibited a certain posture, with her legs retracted and her head tucked under the pronotum.

### Discussion

Male Y frequently *escaped* when under laboratory observation. On the first day of laboratory observation, Male Y *pushed* Male X but did not engage in *pushing* behavior in subsequent treatments. The initial interaction, therefore, appears to have established Male X as the dominant individual, a status that was retained for the duration of the treatments(at least three days). The recurrence of this outcome suggests a mechanism of status recognition and retention that warrants further investigation, because of its centrality to understanding the mating strategy and cognitive abilities of *P. angustipennis*.

Title: “Fight, Retreat, Repeat: Field observation of male-male agonistic behavior in the wood-feeding cockroach, Panesthia angustipennis spadica (Dictyoptera: Blattodea: Blaberidae)”

Authors: Haruka Osaki*, Tomohiro Nakazono, Kiyotaka Yabe, Mamoru Takata, and Aram Mikaelyan

*North Carolina State University;
